# Specificity of Gβ and γ subunits to the SNARE complex both at rest and after α_2a_ adrenergic receptor stimulation

**DOI:** 10.1101/2020.04.29.065136

**Authors:** Yun Young Yim, W. Hayes McDonald, Katherine M. Betke, Ali Kaya, Karren Hyde, Kevin Erreger, Ralf Gilsbach, Lutz Hein, Heidi E. Hamm

## Abstract

Though much is known about the various physiological functions of each GPCR and the specificity of Gα subunits, the specificity of Gβγ activated by a given GPCR and activating each effector in vivo is not known. Previously, we identified different Gβ and Gγ subunits interacting specifically with α_2a_-adrenergic receptors (α_2a_AR). In this study, we examined its in vivo specificity to the soluble NSF attachment proteins (SNARE) complex in adrenergic (auto-α_2a_AR) and non-adrenergic (hetero-α_2a_AR) neurons. We applied a quantitative targeted multiple reaction monitoring proteomic analysis of Gβ and Gγ subunits bound to the SNARE complex, and found only a subset of Gβ and Gγ bound. Without stimulation of auto-α_2a_AR, Gβ_1_ and Gγ_3_ interacted with the SNARE complex. When auto-α_2a_AR were activated, Gβ_1_, Gβ_2_, and Gγ_3_ were found. Further understanding of in vivo Gβγ specificity to its effectors provides new insights into the multiplicity of genes for Gβ and Gγ.

**Summary:** Specific Gβγ dimers interact with the SNARE complex following presynaptic α_2a_AR activation in both adrenergic and non-adrenergic neurons.

## Introduction

Numerous studies have demonstrated that different Gβ and Gγ subunits may selectively interact to form unique Gβγ dimers (*1–3*), which in turn target specific G protein-coupled receptors (GPCRs) and effectors of Gβγ (*1, 3-11*). Although many of these Gβγ-effector interactions and downstream signaling cascades have been well studied in vitro, factors controlling the specificity of Gβγ dimers associated with unique GPCRs or their effectors, as well as the in vivo expression abundance of Gβ and Gγ subunits or Gβγ dimers remain unclear. Only a few in vivo genetic deletion studies and knout-out animal studies have attempted to identify the specific roles of different Gβ and Gγ subunits(*11, 12*). With the recent development of targeted multiple reaction monitoring (MRM) proteomic method to detect each neuronal Gβ and Gγ subunits, our lab has showen the protein abundance and subcellular localization of each neuronal Gβ and Gγ subunits (*13*) and their specificity to α_2a_-adrenergic receptors in adrenergic neurons (auto-α_2a_AR) and non-adrenergic neurons (hetero-α_2a_AR)(*14*). Only a subset of neuronal Gβ and Gγ subunits interact with auto-α_2a_AR and hetero-α_2a_AR. However, it remains unclear how G protein specificity plays out in α_2a_AR-mediated effector interactions.

α_2a_-adrenergic receptors (α_2a_AR) are one of the best studied G_i/o_ coupled GPCRs, given the availability of both knock out (α_2a_AR KO) mice and transgenic mice with tagged receptors (FLAG-α_2a_AR) (*15–19*). Briefly, the activation of α_2a_AR releases GTP-Gα_i/o_ and Gβγ, and these interact with various effectors to initiate the downstream signaling cascades(*11*). FLAG-α_2a_AR transgenic mice express only auto-α_2a_AR under the control of the dopamine-β-hydroxylase (Dbh) promoter while α_2a_AR KO mice are the endogenous knockout of α_2a_AR(*16*). Using these mice, auto-α_2a_AR are found to be important for bradycardia and hypotension while α_2a_AR in hetero-α_2a_AR are involved in anesthetic sparing, hypothermia, analgesia, bradycardia, and hypotension (*16*). However, it is unclear how G protein specificity, especially Gβγ, plays out in α_2a_AR-mediated downstream signaling cascades and underlies such diverse physiological functions. Given the physiological importance of α_2a_AR, and the different roles of auto-and hetero-α_2a_AR, the signaling mechanisms of α_2a_AR in both adrenergic and non-adrenergic neurons, especially in relation to the specificity of Gβγ, need to be further elucidated.

Soluble N-ethylmaleimide-sensitive factor attachment protein receptor (SNARE complex), Gβγ effector, is a key protein involved in synaptic transmission, essential for interneuronal communication in both the nervous and endocrine systems. The neuronal SNARE complex, comprised of SNAP25, syntaxin 1A, and synaptobrevin (VAMP) (*20–24*), brings the vesicle membrane and plasma membrane into close proximity, resulting in vesicle fusion at the plasma membrane (*25*). After membrane fusion, neurotransmitter is released into the synaptic junction, permitting communication between neurons encoding both external and internal sensory inputs, and promoting responsive behaviors in postsynaptic cells (*24*). Thus, it is crucial to understand how synaptic transmission is regulated in healthy subjects, and mechanisms of dysfunction in neurological diseases (*26–28*).

The activation of α_2a_AR results in the inhibition of synaptic transmission via the Gβγ subunit in both adrenergic (auto-α_2a_AR) and non-adrenergic neurons (hetero-α_2a_AR) such as glutamatergic, GABAergic, serotonergic and dopaminergic neurons (*29–32*). In particular, numerous studies have reported that Gβγ subunits released from activated α_2a_AR reduce presynaptic Ca^2+^ influx through the inhibition of voltage-gated calcium channels (VDCC) (*33–35*) and inhibit exocytosis through direct interaction with the SNARE complex (*36–38*). Furthermore, the direct interaction between Gβy and the SNARE complex upon α_2a_AR activation is a primary mechanism to inhibit exocytosis of norepinephrine-containing vesicles in the central amygdala (*38*). Delaney *et al*. show that inhibition of norepinephrine release, where high abundance of presynaptic α_2a_AR are present, is mediated by a G_i/o_-coupled receptor, and that Gβγ dimers released from active α_2a_AR directly interact with the SNARE complex downstream of calcium entry to inhibit norepinephrine release (*38*). In addition, our knout-out study with SNAP25Δ3 mice no longer shows the inhibition of adrenergic signaling in the BNST, a key area for the integration of stress signals (*39*). The loss of 3 C-terminal residues in SNAP25 causes a loss of Gβγ‘s ability to regulate α_2a_AR mediated inhibition of synaptic transmission. Although the Gβγ-SNARE complex interaction has been well-characterized during α_2a_AR-mediated inhibition of synaptic transmission, the specificity of Gβγ dimer isoforms for the SNARE complex is not known.

Together, we use various biochemical techniques and tools such as coimmunoprecipitation (coIP), quantitative MRM proteomic method, and transgenic animals to understand the specificities of Gβ and Gγ subunits to the SNARE complex. Here, we evaluated which Gβγ dimers released from epinephrine (epi)-activated α_2a_AR interact with SNARE complex to regulate synaptic transmission. We discovered that only a subset of Gβ and Gγ subunits released from activated α_2a_AR interact with the SNARE complex. Further understanding of α_2a_AR-mediated Gβγ subunit specific interaction with the SNARE complex may lead to the development of new therapeutic targets that complement or contrast with currently available α_2a_AR-targeted drugs.

## Results

### Gβγ - SNARE complex interaction

To study Gβγ specificity for the SNARE complex upon activation of α_2a_AR, we used synaptosomes collected from the brains of wildtype, α_2a_AR KO, and FLAG-α_2a_AR mice. Using a SNAP25 antibody (S9684) and DSP, a lipid-soluble thiol cleavable crosslinker, Gβγ and the SNARE complex were co-immunoprecipitated (coIP) and detected by Western blot (Fig. 1A). We showed that the SNAP25 (S9684) antibody selectively recognizes SNAP25 (fig.S1). To further validate that we were indeed coIPing the ternary SNARE complex, we detected Syntaxin1 in addition to SNAP25 from these coIP samples (fig. S2). The Gβγ and SNARE complex interaction was found in both unstimulated and stimulated coIP samples, but greater amounts were present in epi-stimulated coIP samples of wildtype and FLAG-α_2a_-AR mice (Fig.1B to E). We know from a number of studies (*40, 41*) that there is a low affinity for interaction between Gβγ and the SNARE complex. However, the interaction is specific, as little non-specific binding was observed in IgG controls. Quantitative western blot analysis estimated approximately 1μg Gβγ was pulled down with SNAP25 per half brain (fig. S3). Together, these results suggest that this method is suitable to determine which Gβγ subunits associate with the SNARE complex in vivo.

**Figure 1.**
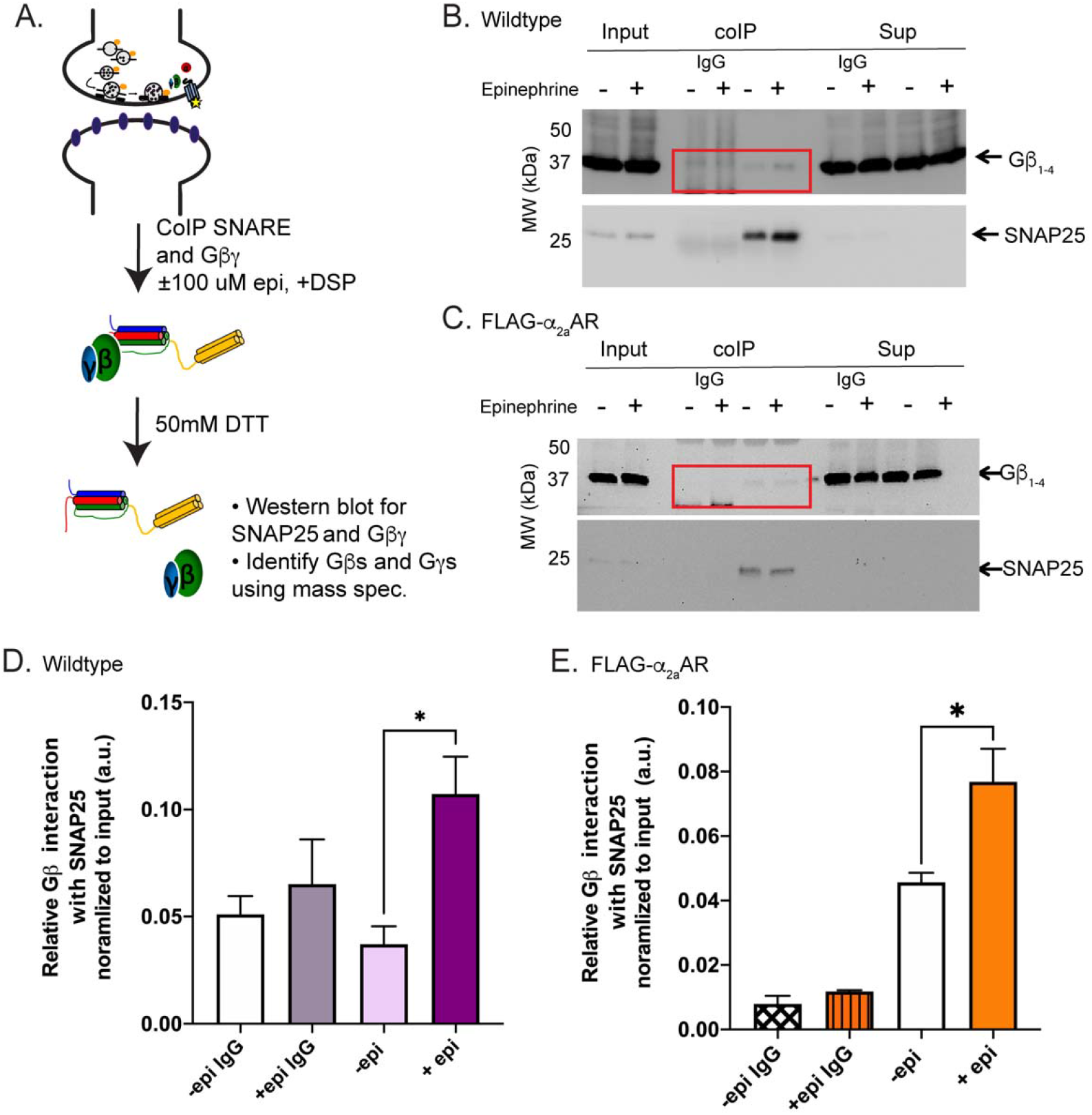
Co-immunoprecipitation of the SNARE complex and Gβγ. Coimmunoprecipitation (coIP) experimental protocol (**A**), and representative western blot of coimmunoprecipitation of SNAP25 and Gβγ in wildtype (**B**) or FLAG-α_2a_AR (**C**) mice following the stimulation of synaptosomes with (+, 100μM epi) or without epi (-). DSP, a lipid-soluble thiol cleavable crosslinker, was added to ensure the receptor and Gβγ remained intact during co-IP experiments. We detected Gβγ in the SNAP25 pulldown but not the IgG control pulldown regardless of receptor stimulation (**B** and **C**, red box). Quantified Gβ intensities were normalized to that of input and presented as relative fold change in wildtype (**D**) or FLAG-α_2a_AR (**E**) mice (N=3). Data are presented as mean ± SEM. One-way ANOVA* p<0.05, **p<0.01, *Post hoc* analysis by Tukey’s multiple comparison. coIP: co-immunoprecipitated fraction; Sup: depleted supernatant

### Gβ_1_, Gβ_2_, Gβ_4_, Gγ_2_, Gγ_3_, and Gγ_4_ interact specifically with the SNARE complex upon epinephrine stimulation

To examine which neuronal Gβ and Gγ subunits interact with the SNARE complex upon activation of α_2a_-AR, we applied a quantitative MRM method (*14*) to TCA precipitated and trypsin digested SNAP25 coIP samples of wildtype (WT), α_2a_AR KO, and FLAG-α_2a_AR mouse synaptosomes. Previously, only a subset of neuronal Gβ and Gγ subunits are selectively associated with α_2a_AR(*14*). WT samples were used to detect the Gβγ-SNARE complex interactions mediated by epi stimulation, including both auto- and hetero-α_2a_AR, while α_2a_AR KO samples were used to detect non α_2a_AR-mediated interactions only. Lastly, FLAG-α_2a_AR samples were used to detect presynaptic auto-α_2a_AR-mediated Gβγ-SNARE complex interactions. We used unstimulated WT samples (WT no epi), α_2a_AR KO (α_2a_AR KO no epi), and FLAG-a_2a_AR (FLAG-α_2a_ARs no epi) as controls to detect basal Gβγ-SNARE complex interactions. By comparing the amount of Gβ and Gγ subunits detected per mg of lysate between stimulated WT (WT +epi), α_2a_AR KO (α_2a_AR KO +epi), and FLAG-α_2a_AR (FLAG-α_2a_AR +epi), we detected not only auto-receptor mediated Gβγ-SNARE complex interactions, but also quantify those mediated by non-α_2a_AR and estimated heteroreceptor interactions.

To identify nonspecific interactions of Gβ and Gγ subunits, we used IgG alone controls (without primary antibody). We compared the amount of Gβ subunit detected per mg of lysate between WT IgG +epi and WT +epi samples and found Gβ_1_, Gβ_2_, and Gβ_4_ significantly interacted with the SNARE complex upon α_2a_AR stimulation (Fig. 2). As expected from previous studies (*13, 14*), Gβ_1_ was detected at a significantly higher concentration than Gβ_2_ and Gβ_4_ subunits. Moreover, we did not identify interaction between Gβ_5_ and the SNARE complex following α_2a_AR activation.

**Figure 2.**
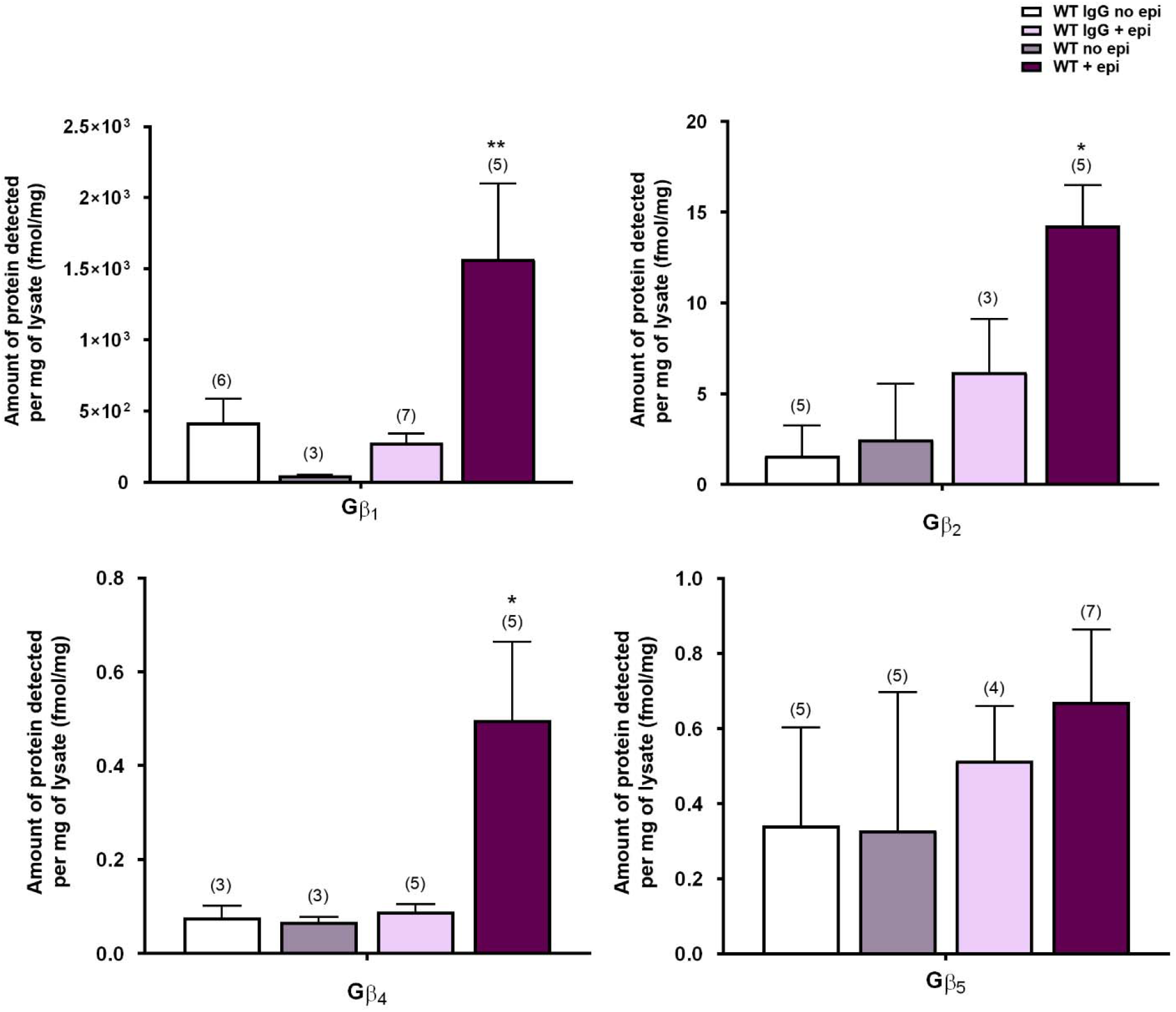
Gβ specificity to the SNARE complex upon α_2a_AR activation. Quantification of Gβ subunits interacting with the SNARE complex in wildtype synaptosomes (N=8, unless otherwise noted). Gβ_1_, Gβ_2_, and Gβ_4_ interact specifically with the SNARE complex upon α_2a_AR activation with epi. Data are presented as mean ± SEM. One-way ANOVA* p<0.05, **p<0.01, *Post hoc* analysis by Tukey’s multiple comparison.

Next, we identified the Gγ subunits that interact with the SNARE complex following receptor activation. We found enrichment of the Gγ_2_, Gγ_3_, and Gγ_4_ subunits with the SNARE complex upon epi stimulation (Fig.3). Gγ_3_ was the most abundant followed by Gγ_2_ and Gγ_4_, respectively. We detected a similar abundance of Gγ_2_ and Gγ_4_ associated with the SNARE complex. We were not able to detect an interaction between Gγ_12_ and the SNARE complex (Fig. 3). Although Gγ_12_ is specifically bound to α_2a_AR (*14*), it may not interact with the SNARE complex but rather with other Gβγ effectors such as VDCCs (see Discussion). Even though we are not detecting specific Gβγ dimers in this study, according to our data (Fig. 2 and 3), we detect only a subset of possible Gβγ dimers and can postulate 9 different combinations of Gβγ dimers (Gβ_1_γ_2_, Gβ_1_γ_3_, Gβ_1_γ_4_, Gβ_2_γ_2_, Gβ_2_γ_3_, Gβ_2_γ_4_, Gβ_4_γ_2_, Gβ_4_γ_3_, and Gβ_4_γ_4_) which might interact with the SNARE complex upon epi stimulation. These dimers may occur in the brain and play roles in the α_2a_AR mediated Gβγ-SNARE complex interactions. Based on the detected abundances of Gβ_1_ and Gγ_3_, we suggest that Gβ_1_γ_3_ is the most likely dimer to interact with the SNARE complex following receptor stimulation. Further biochemical analysis will be needed to validate the presence of these Gβγ dimers and their specificities for the SNARE complex upon α_2a_AR activation in both adrenergic and non-adrenergic neurons.

**Figure 3.**
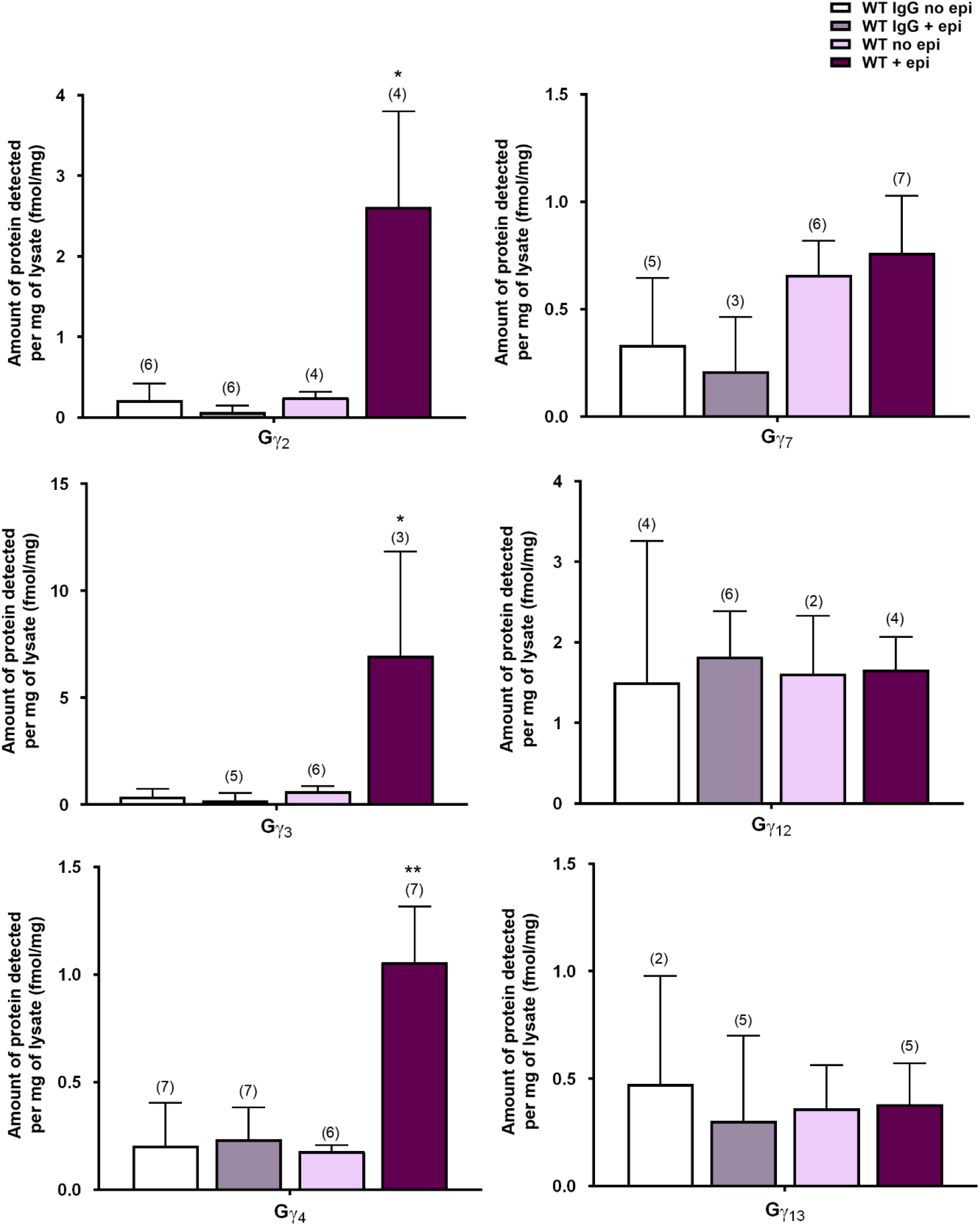
Gγ specificity to the SNARE complex upon α_2a_AR activation. Quantification of Gγ subunits interacting with the SNARE complex in wildtype synaptosomes (N=8,unless otherwise noted). Gγ_2_, Gγ_3_, and Gγ_4_ may specifically interact with the SNARE complex upon α_2a_AR activation. Data are presented as mean ± SEM. One-way ANOVA* p<0.05, **p<0.01, *Post hoc* analysis by Tukey’s multiple comparison.

### Basal interaction of Gβγ and the SNARE complex

We showed in Fig 1B and C that Gβγ and the SNARE complex interact basally in unstimulated coIPed samples of wildtype and FLAG-α_2a_-AR mice. To understand the basal Gβγ-SNARE complex interaction in the absence of epi stimulation, we compared IgG control to no epi coIPed samples from α_2a_AR KO and FLAG-α_2a_AR synaptosomes using MRM mass spectrometry. α_2a_AR KO samples were used to identify non-α_2a_AR-mediated basal interactions while FLAG-α_2a_AR samples were used to detect auto-α_2a_AR-mediated Gβγ-SNARE complex interactions. Here, we detected all neuronal Gβ subunits but found only Gβ_1_ significantly interacting with the SNARE complex in both α_2a_AR KO and FLAG-α_2a_AR no epi conditions compared to that of the IgG no epi condition (Fig.4). This suggests that there is a basal interaction of Gβ_1_ with the SNARE complex. The abundance of Gβ_1_ between these two conditions was not different, suggesting that the basal interaction of Gβ_1_ and the SNARE complex may not be α_2a_AR mediated. Similarly, we detected Gγ_3_ interaction with the SNARE complex in the unstimulated conditions of both genotypes compared to IgG control (Figure 5). Thus we suggest that the Gβ_1_γ_3_ dimer is a basal, non-α_2a_AR mediated interaction. Various GPCRs have been implicated to mediate presynaptic inhibition through this mechanism; therefore, other endogenous GPCRs present in the synaptosomes could be causing this effect, or alternatively, since no agonists of other GPCRs were present in the experiments, it may be a basal interaction with SNARE complex

**Figure 4.**
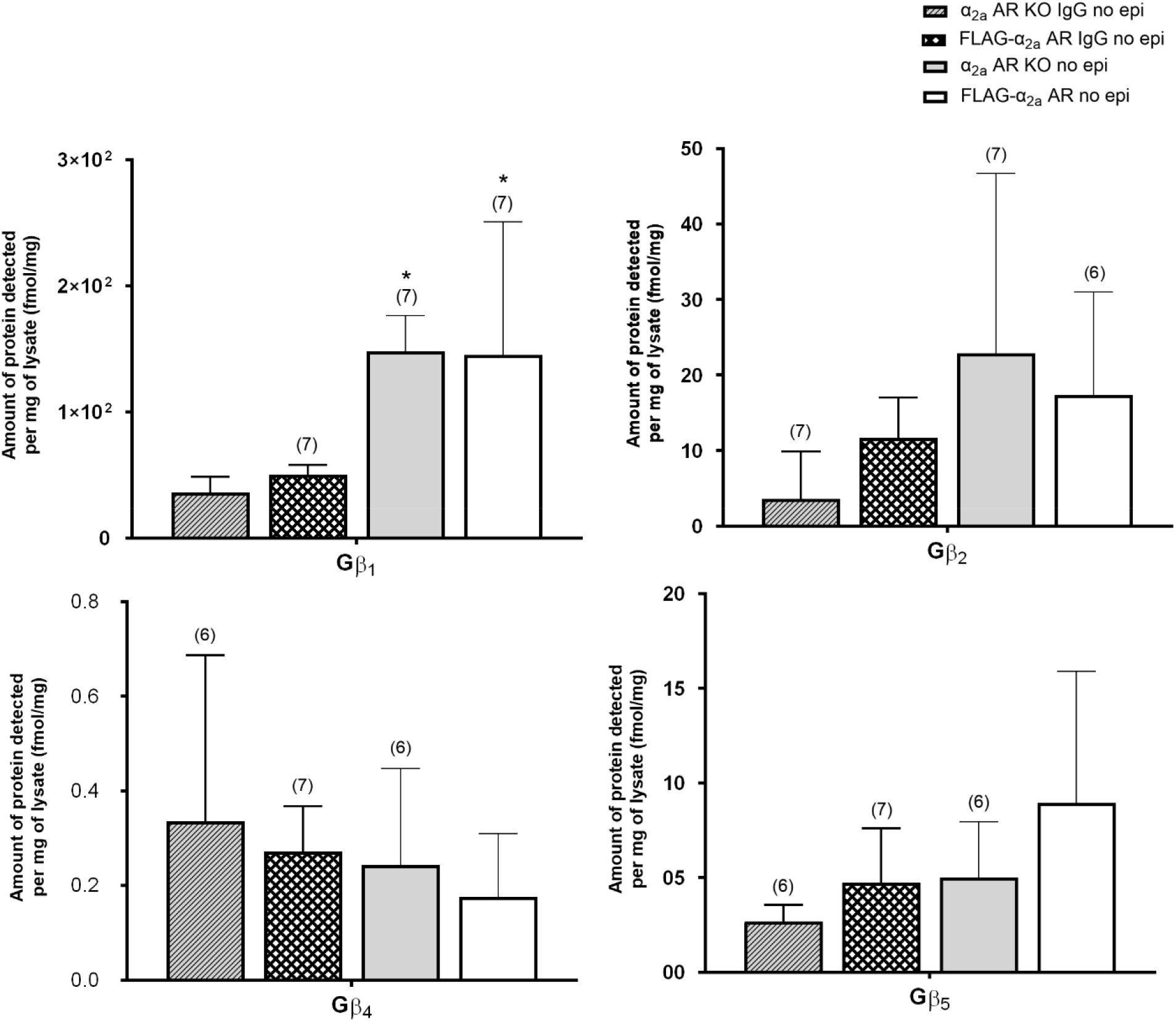
Gβ subunits interacting basally with the SNARE complex. Quantification of Gβ subunits interacting with the SNARE complex basally (N=8, unless otherwise noted). Gβ_1_, but not Gβ_2_, specifically interacting with the SNARE complex in both α_2a_AR KO and FLAG-α_2a_AR no epi conditions compared to that of IgG no epi. Data are presented as mean ± SEM. One-way ANOVA* p<0.05, **p<0.01, *Post hoc* analysis by Tukey’s multiple comparison.

**Figure 5.**
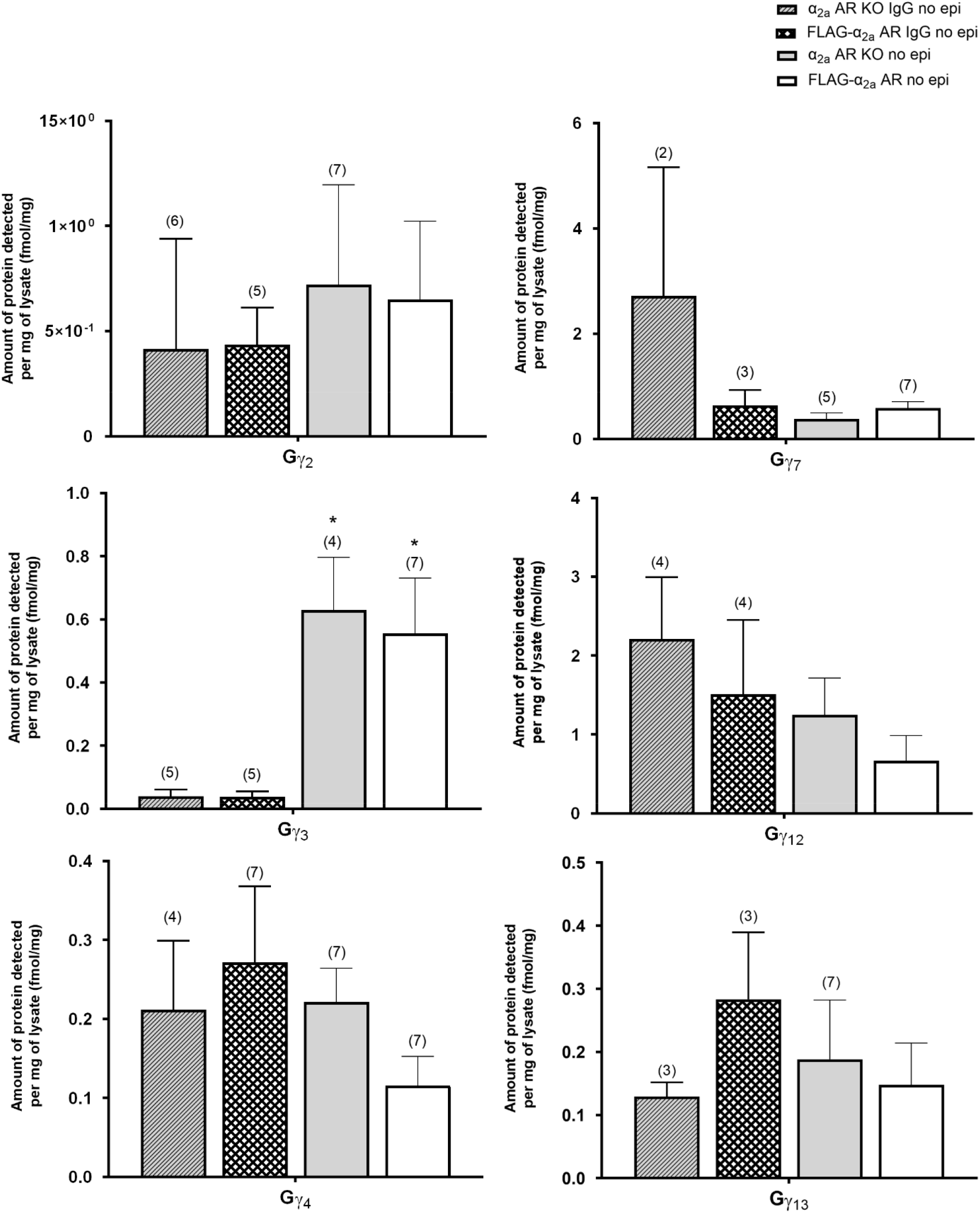
Gγ subunits interacting basally with the SNARE complex. Quantification of Gγ subunits interacting with the SNARE complex basally (N=8, unless otherwise noted). Gγ_3_ specifically interacted with the SNARE complex in both α_2a_AR KO and FLAG-α_2a_AR no epi conditions compared to that of IgG no epi. Data are presented as mean ± SEM. One-way ANOVA* p<0.05, **p<0.01, *Post hoc* analysis by Tukey’s multiple comparison.

### Gβ_1_, Gβ_2_, and Gγ_3_ interact specifically with the SNARE complex upon auto-α_2a_ARs activation

As previously described, adrenergic auto-α_2a_AR inhibits presynaptic neurotransmitter release through the interaction between Gβγ and the SNARE complex (*38*). To test whether auto-α_2a_ARs and presynaptic hetero-α_2a_AR utilize different Gβγ dimers upon activation, we coIPed samples from α_2a_AR KO and FLAG-α_2a_AR synaptosomes and again applied the quantitative MRM method to understand auto-α_2a_ARs mediated Gβγ-SNARE complex interactions. α_2a_AR KO (no epi) and FLAG-α_2a_AR (no epi) samples were used as controls to identify non-auto-α_2a_AR mediated interactions, and α_2a_AR KO +epi samples were used to detect non-α_2a_AR-mediated Gβγ-SNARE complex interactions. By comparing the result from α_2a_AR KO, FLAG-α_2a_AR, and wildtype, we can propose the specificity of Gβ and Gγ interactions with the SNARE complex mediated by hetero-α_2a_AR.

Here, we were able to detect a subset of Gβ and Gγ subunit interactions with the SNARE complex upon auto-α_2a_AR activation. When we compared the results from α_2a_AR KO +epi and FLAG-α_2a_AR +epi synaptosomes, we identified that Gβ_1_ and Gβ_2_, but not Gβ_4_, enrich with the SNARE complex following auto-α_2a_AR activation (Fig.6). We also detected significantly more Gβ_1_ than Gβ_2_ (Fig.6). Moreover, we detected a ~ 10-fold increased interaction of Gβ_1_ and the SNARE complex upon epi stimulation compared to the basal state (compare Fig. 4 and 6).

**Figure 6.**
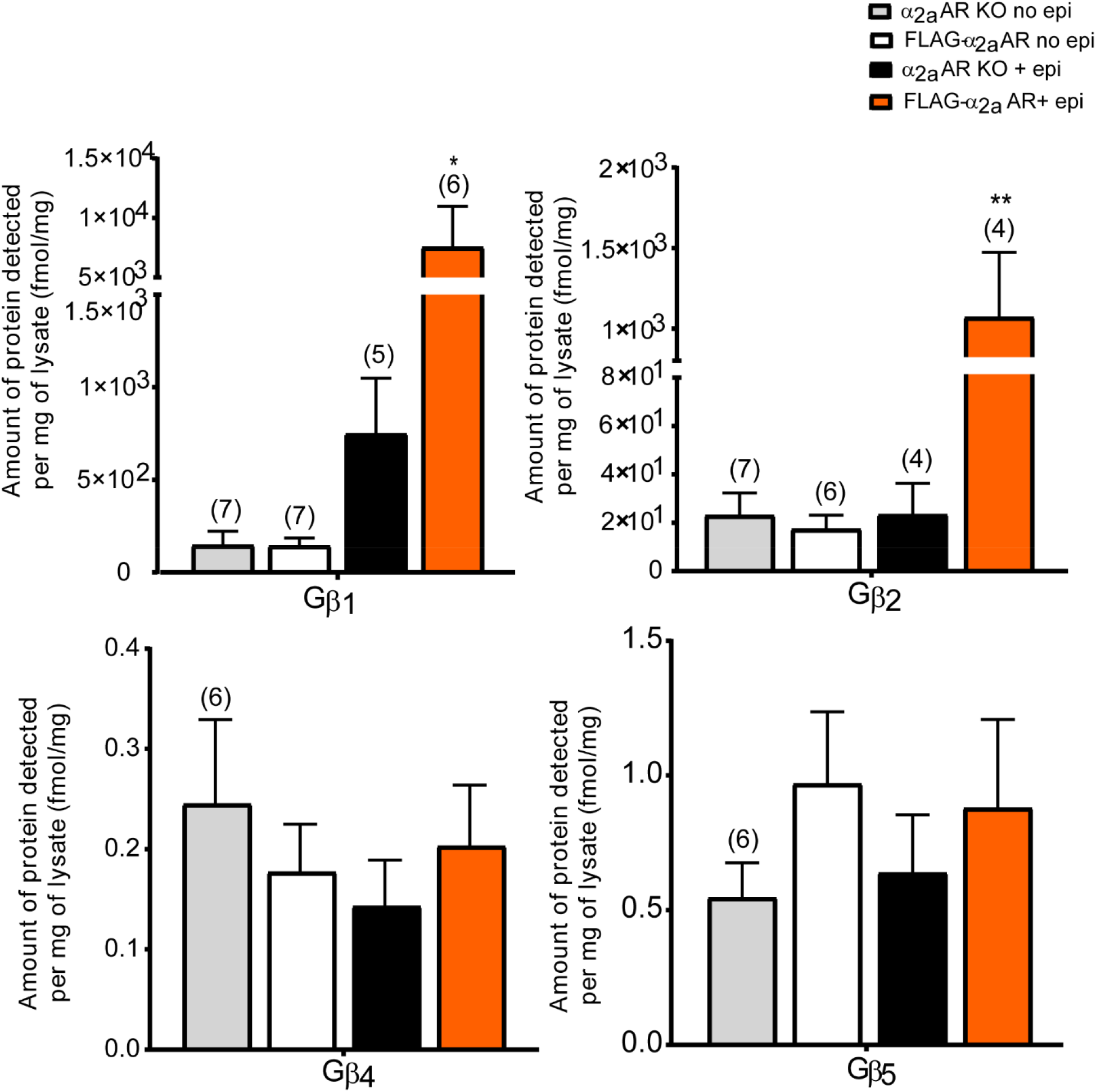
Gβ specificity to the SNARE complex upon auto-α_2a_ARs activation. Quantification of Gβ subunits interacting with the SNARE complex in α_2a_AR KO and FLAG-α_2a_AR synaptosomes (N=8, unless otherwise noted). Gβ_1_ and Gβ_2_ specifically interact with the SNARE complex upon auto-α_2a_AR activation. Data are presented as mean ± SEM. One-way ANOVA* p<0.05, **p<0.01, *Post hoc* analysis by Tukey’s multiple comparison.

Among the 3 Gγ subunits enriched with the SNARE complex upon stimulation of WT containing auto- and hetero-α_2a_AR and non-α_2a_AR (Fig. 3), the enrichment of Gγ_3_ with the SNARE complex was detected following auto-α_2a_AR activation (Fig. 7). Basally, without epi stimulation, Gγ_3_ was found to interact with the SNARE complex (Fig 5). These findings suggest that Gβ_1_γ_3_ may be basally interacting with the SNARE complex and not α_2a_ARs mediated (Fig. 4 and 5), whereas Gβ_2_γ_3_ may interact with the SNARE complex only after epi stimulation of auto-α_2a_AR.

**Figure 7.**
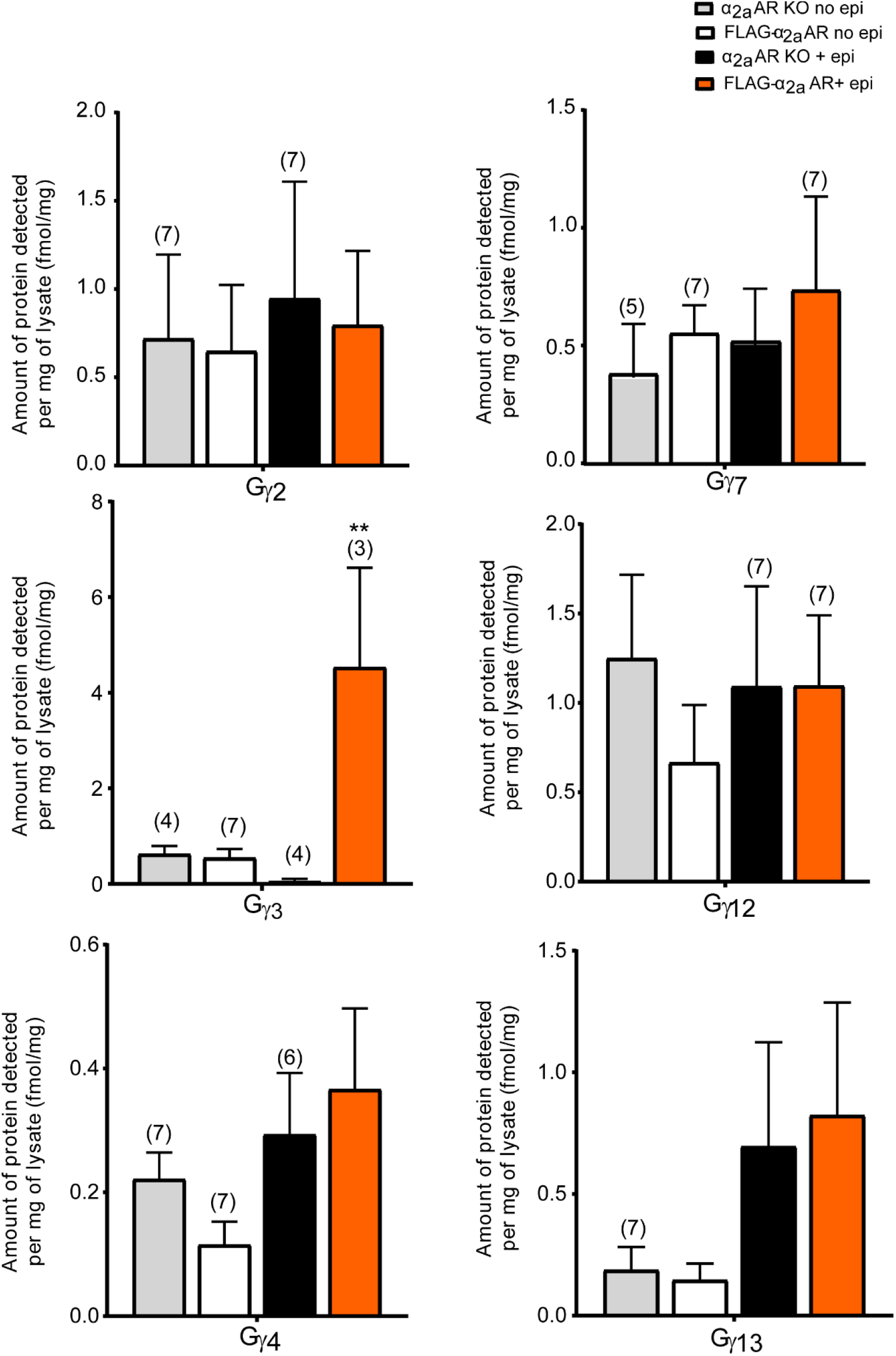
Gγ subunit specificity to auto-α_2a_ adrenergic receptors. Quantification of Gγ subunits interacting with the SNARE complex in α_2a_ARs KO and FLAG-α_2a_AR synaptosomes (N=8, unless otherwise noted). Gγ_3_ specifically interacts with SNARE upon auto-α_2a_AR activation. Data are presented as mean ± SEM. One-way ANOVA* p<0.05, **p<0.01, *Post hoc* analysis by Tukey’s multiple comparison.

**Figure 8.**
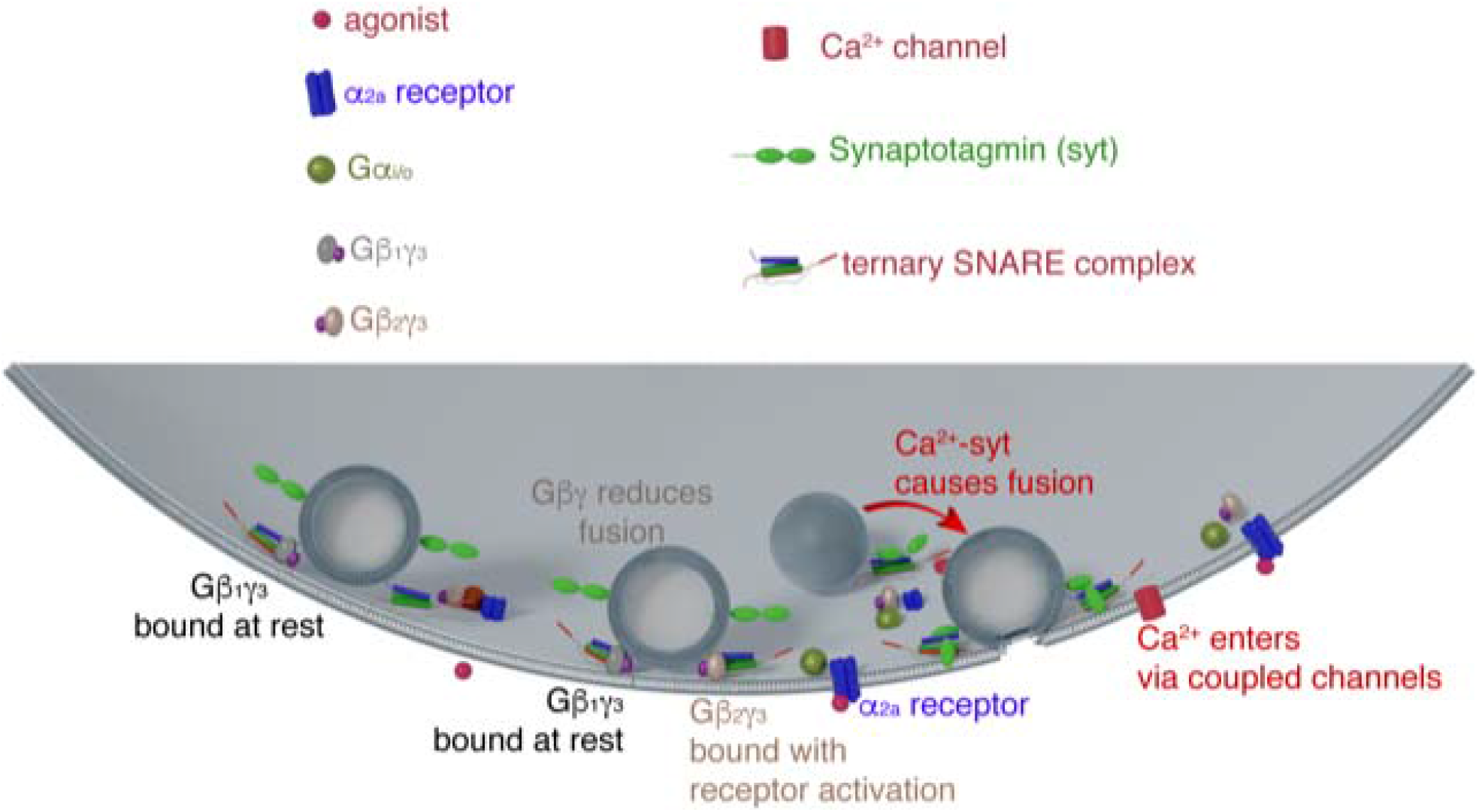
Summary Figure. While the Gβγ-SNARE complex interaction may exist basally (ex. Gβ_1_γ_3_), only a subset of Gβγ dimers made with Gβ_1_ Gβ_2_, Gβ_4_, Gγ_2_, Gγ_3_, and Gγ_4_ (ex. Gβ_2_γ_3_) specifically interact with the SNARE complex upon presynaptic α_2a_AR activation in both adrenergic and non-adrenergic neurons.

## Discussion

Numerous in vitro studies have studied the effectors of Gβγ and their downstream signaling cascades(*1-4, 42*); however, only a few in vivo genetic deletion studies and knout-out animals studies have examined the specific roles of different Gβ and Gγ subunits (*11, 12*). It remains unclear how particular Gβγ dimers form or how they localize to particular sites in vivo. In this paper, we addressed the in vivo specificity of Gβγ subunit interaction with the SNARE complex. We determined a surprising specificity in which Gβ and γ subunits associate with the SNARE complex, including the interesting finding that some Gβγs interact with the SNARE complex regardless of receptor activation. Only a subset of neuronal Gβγ dimers—Gβ_1_, Gβ_2_, Gβ_4_, Gγ_2_, Gγ_3_, and Gγ_4_—interact with the SNARE complex (Fig. 2 and 3) following epi stimulation in WT mice. Upon the activation of auto-α_2a_AR, only Gβ_1_, Gβ_2_, and Gγ_3_ interacted with the SNARE complex (Fig. 6 and 7). From 9 possible Gβγ dimers, auto-α_2a_AR may utilize only two Gβγ dimers, Gβ_1_γ_3_ and Gβ_2_γ_3_, to inhibit neurotransmitter release through the Gβγ-SNARE complex interaction, In addition, comparing WT and auto-α_2a_AR samples (Fig.2 and 6), we found that Gβ_4_γ_2_ and Gβ_4_γ_4_ may be specifically utilized by presynaptic hetero-α_2a_AR. Thus, this study provides a window into the specificity of Gβγ dimer function in vivo.

In both Western blot and quantitative MRM study in FLAG-α_2a_AR animals, we detected Gβγ-SNARE complex interactions in the basal state, without α_2a_AR stimulation (Fig.1C, 4, and 5). Because we showed previously that the α_2a_AR does not basally interact with Gβγ (*14*) and no basal Gβγ interaction to the SNARE complex was detected in the quantitative MRM study of wildtype animals (Fig. 2), we were surprised to see a basal interaction of Gβ_1_γ_3_ with the SNARE complex (Fig.4 and 5). With the limited tools, we cannot evaluate and understand the observed difference of basal interaction in wildtype and FLAG-α_2a_AR animals However, the ~4-fold increase of abundance in Gβ and ~20-fold increase of abundance in Gγ observed in FLAG-α_2a_AR animals raise an interacting hypothesis. The change in Gβγ associated to the SNARE complex may occur upon the activation of auto-α_2a_AR. Further studies are necessary to understand the basal interaction of Gβ_1_γ_3_ with the SNARE complex, especially in the wildtype animal Gβ_2_γ_3_ appears only associated with the SNARE complex upon auto-α_2a_AR activation.

Gβ_1_ specifically interacts with the SNARE complex not only in the basal state and but also upon epi stimulation in FLAG-α_2a_AR mice (Fig.4 and 6). The abundance of Gβ_1_ interacting with the SNARE complex in WT samples (Fig. 2) was expected as Gβ_1_ is the most abundant Gβ subunit in whole synaptosome as well as at both pre-and post-synaptic fractions(*13*). However, an increase of abundance in Gβ_1_ (~10 fold) interacting with the SNARE complex upon auto-α_2a_AR activation was not expected (Fig.6). This suggests that the basally bound Gβ_1_ may have a unique role in basal modulation of exocytosis. So far, the role of this basal Gβγ association with the SNARE complex is not known, but it is well-known that basal exocytotic events (measured electrophysiologically as mini EPSCs) can be regulated by G_i/o_ GPCR agonists(*43, 44*).

Combining this study of the Gβγ specificity to the SNARE complex with our previous studies of the presynaptic abundance specificity of Gβγ subunits for α_2a_AR(*13, 14*), we hypothesize that the Gβγ subunit specificity may be encoded by an association of a particular GPCR with a subset of Gβγs, –the abundance of each neuronal Gβ and Gγ subunit, and the distance to their effectors at the presynaptic terminal. In our previous receptor study comparing the protein abundance detected with α_2a_ receptors, we speculated that Gγ_2_, Gγ_3_, and Gγ_4_ were associated with auto-α_2a_AR while Gγ_12_ is specific for hetero-α_2a_AR (*14*). Given our current understanding with this study on the Gβ and Gγ specificity to the SNARE complex, we can more accurately show that Gγ_2_ and Gγ_4_ are involved in hetero-α_2a_AR-mediated Gβγ-SNARE complex interaction, with no evidence for Gγ_12_ association to the SNARE complex. Gγ_12_ associated with hetero-α_2a_AR may be associated with another Gβγ effector.

The Gγ subunits, more than Gβ, play key roles in Gβγ specificity. Previously, knockdown and knock-out studies showed that particular Gγ subunits have distinct physiological importance in biological systems (*11, 12*). To date, only 4 out of 12 Gγ subunits have been studied using knockdown or knock-out models. Mainly expressed in retina, Gγ_1_ is involved in photoreceptor degeneration. Gγ_2_ is involved in nociception and angiogenesis. Gγ_3_ is involved in increased susceptibility to seizures and resistance to diet induced obesity while Gγ_7_ affects the startle response(*4, 7*). Unique physiological phenotypes of each Gγ subunit suggest a great deal of specificity in Gβγ dimerization and signaling(*45–47*). The main molecular mechanisms behind binding selectivity between various Gβγs and their effectors are still unknown. However, ongoing investigations of interactions between each Gγ and t-SNARE are suggesting the importance of charged residues during this process(*48*).

Moreover, the fast rate of Gβγ-SNARE complex-mediated inhibition(*44*) suggests that Gβγ, the SNARE complex, and other synaptic proteins may be associated by scaffolding interactions prior to receptor activation, to facilitate Gβγ-SNARE complex interaction and mediate Gβγ specificity. Previously, we identified two Gβγ binding sites on vesicle proximal and distal regions of SNAP25 (*41, 49, 50*), far enough apart to not both be occupied by a single Gβγ. However, the physiological and biochemical importance of these interaction sites still remains unclear. The two sites could underlie interaction via the SNARE complex with two different effectors (*51, 52*) or the amino terminal region may be a Gβγ binding (or “holding”) site that can hand Gβγ off to the physiologically important carboxy-terminal interaction site (*50*). Using SNAP25Δ3 mice with a truncated C-terminus of SNAP25, we see a two-fold reduction in Gβγ-SNARE complex interaction, as well as an increase in stress-induced hyperthermia, defective spatial learning, impaired gait, supraspinal nociception (*39*), enhancement of long-term potentiation at Schaffer collateral-CA1 synapse (*53*), and resistance to diet-induced obesity(*54*). It will be interesting to determine the stoichiometry of Gβγ binding to the SNARE complexes, and whether identical Gβγ dimers bind to both amino- and carboxy-terminal sites of the SNARE complexes. Detailed quantitative analysis in combination with SNAP25 mutagenesis studies will be necessary to answer these questions.

The specificity of Gβγ dimers to the SNARE complex is not fully understood, but its importance and the relationship of GPCRs to this interaction have been well-studied. Hamid *et al*. demonstrated that 5HT_1b_ and GABA_B_ receptors inhibit exocytotic release at the same synaptic terminal through different mechanisms (*43*), with 5HT_1b_ receptor-mediated inhibition of exocytosis using the Gβγ-SNARE complex interaction, while GABA_B_ receptor-mediated inhibition works through Gβγ-VDCC interaction and decreased calcium entry. The authors suggested that this difference may depend on the N-terminal SNAP25 residues. The association of specific Gβγ subunits to GPCRs, such as 5HT_1b_ and GABA_B_, and to the SNARE complex may add a further layer of regulation of Gβγ-effector selectivity. Future studies to understand this specificity will clarify Gβγ isoform specificity important for both receptor and effector binding.

## Materials and Methods

### Animals

Adult, male FLAG-tagged alpha2a adrenergic receptors (α_2a_-AR), α_2a_-AR knock-out (KO), and wildtype mice (*55*) were decapitated and brain tissues were immediately homogenized to produce crude synaptosomes as described below. To minimize post-mortem differences, all tissues were processed in parallel. All animal handling and procedures were conducted in accordance with the Care and Use of Laboratory Animals of the National Institutes of Health and approved by the Vanderbilt Institutional Animal Care and Use Committee.

### Drugs

Epinephrine (catalog E4642), prazosin (catalog P7791), and propranolol (catalog P0884) were purchased from Sigma-Aldrich.

### Antibodies

For immunoprecipitation, rabbit anti-SNAP25 (Sigma, S9684) and rabbit ChromePure IgG (Jaskson Immuno Research, 011-030-003) were used. For the western blot analysis, mouse anti-SNAP25 (Santa Cruz, sc-376713, 1:500) and rabbit anti-Gβ (Santa Cruz, sc-378, 1:10,000 and 1:5000) were used. HRP-conjugated secondary antibodies were obtained from Perkin-Elmer and Jackson Immunoresearch and used at the following dilutions: goat antimouse (1:10,000), and mouse anti-rabbit light chain specific (1:7,500).

### Synaptosome preparation, stimulation, and lysate protocol

Crude synaptosomes were isolated from mouse brain tissue, as described previously (*14, 56-58*) in the presence of 1mM ethylenediaminetetraacetic acid, 1uM propranolol, 1uM prazosin, and 2mM 3,3’-dithiobis [sulfosuccinimydlypropinate], the lipid soluble, thiol cleavable crosslinker (Pierce, 22585) and stimulated with 100μM epinephrine (epi). This mimics the local synaptic concentration of epi and it is a commonly used concentration in α_2a_AR studies(*59–61*). As epi is not an α2aAR selective agonist, we also used prazosin and propranolol to prevent off-target effects from non-α2aAR adrenergic receptors. Samples were frozen in liquid nitrogen and stored at −80°C.

### Co-immunoprecipitation

Similar to the co-immunoprecipitation (coIP) of α_2a_AR and Gβγ (*14*), we coIPed the SNARE complex and Gβγ from the precleared lysates (Fig.1A). Precleared lysates were aliquoted 400ul per tube. Instead of anti-HA or FLAG antibodies, they were incubated with an anti-SNAP25 coIP antibody. Because homozygous SNAP25 knock-out mice exhibit neonatal lethally(*62*), we used rabbit IgG as a control.

### Elution and trichloroacetic acid (TCA) precipitation of coIPs

Samples (approximately 11 coIPs per condition per genotype) were eluted using 100mM glycine (pH 2.5). 40μL of glycine was added to coIP beads and placed on a room temperature vortex shaker at medium speed for 8 mins and centrifuged at 5,000 x g for 2mins to pellet beads. Elutants were transferred to clean 1.5mL Eppendorf tubes. The elution was repeated a second time, and the elutants were pooled together. For western blot analysis, one tube per condition per genotype was saved separately Remaining elutants were pooled per genotype per condition and TCA precipitated to concentrate the Gβ and Gγ subunits for MRM (*14*). All samples were stored at −80°C freezer for the western blot or MRM analysis.

### Immunoblot Analysis

The analysis was done as described previously (*14*). We used 15%, instead of 10% SDS-PAGE gel.

### Heavy labeled peptide cocktail

A heavy labeled peptide cocktail was made as described previously (*13*).

### Quantitative MRM of Gβ and Gγ subunits

As described previously (*14*), coIP samples containing Gβ and Gγ subunits were separated by SDS-PAGE, in-gel trypsin digested, and analyzed by scheduled multiple reaction monitoring (MRM) using a TSQ vantage triple quadrupole mass spectrometer (Thermo Scientific) coupled directly to a nanoAcquity liquid chromatography system (Waters).

### Statistical analysis

One-way analysis of variance (ANOVA) with a Tukey post hoc test was used to account for differences in protein abundance of Gβ and Gγ subunits and for the quantification of western blots (* p<0.05, ** p<0.01, *** p<0.001). All statistical tests were performed using GraphPad Prism v.9.0 for Windows, (GraphPad Software, La Jolla, California, USA, www.graphpad.com).

## Supplementary Materials

Figure 1. Selectivity of SNAP25 IP Antibody

Figure 2. Western blot analysis of SNAP25 coIP.

Figure 3. Quantification of Gβ precipitated with SNAP25.

## Acknowledgments

We thank the proteomics core of the Mass Spectrometry research Center for advice and technical assistance and Drs. Boris Alten and Ege Kavalali for sharing SNAP25 WT and KO hippocampal culture with us. We would also like to thank Dr. Simon Alford for the illustration of summary figure. Lastly, we would like to thank Dr. Analisa Thompson Gray for reviewing and confirming the statistical tests we used for the analysis.

## Funding

This work was supported by the National Institutes of Health (EY10291, MH101679, MH081917, and T32 GM07628).

## Authors Contributions

Y.Y., W.H.M., K.M.B., and H.E.H. participated in research design and preliminary study. Y.Y. and K.E. conducted experiments and data analysis. W.H.M. ran mass spectrometry and helped with data analysis. K.H., A.K., R.G., and L.H. contributed in mouse breeding and sampling. Y.Y. A.K., K.M.B. and H.E.H. wrote or contributed to the writing of the manuscript. All authors reviewed the results and approved the final version of the manuscript.

## Competing Interests

All authors declare that they have no competing interests.

## Data Availability

All data supporting the findings of this study are available from the lead author Heidi E. Hamm upon reasonable request.

## Notes

### Competing Interest Statement

The authors have declared no competing interest.

