## Supplemental material for "Specificity of Gβ and γ subunits to the SNARE complex both at rest and after α_2a_ adrenergic receptor stimulation"

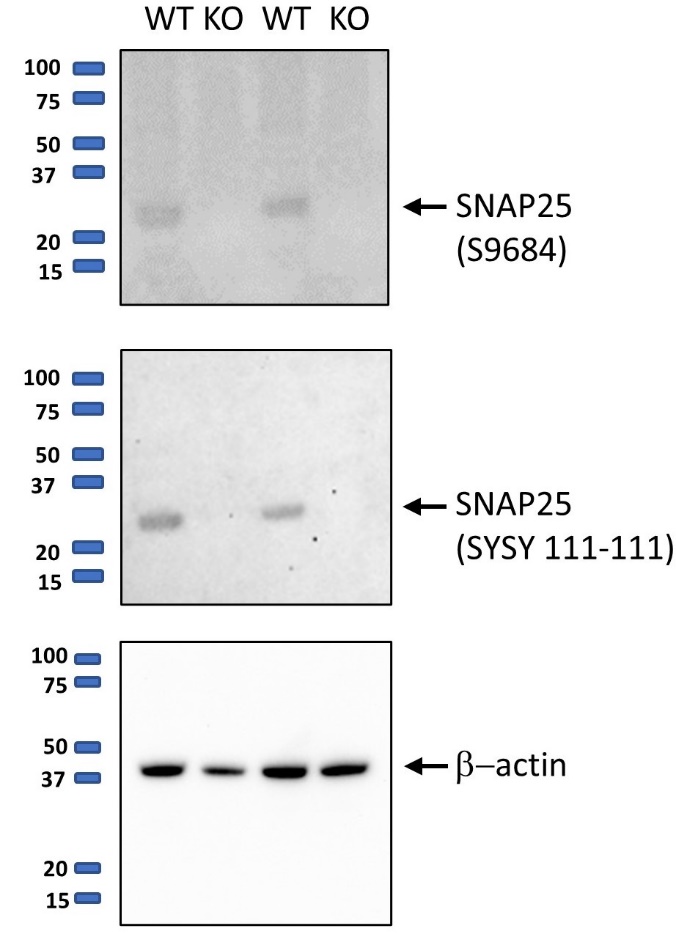

**Supplementary figure 1. Selectivity of SNAP25 IP Antibody.**  To test the selectivity of Sigma SNAP25 IP (S9684) antibody, we carried out Western blot analysis against SNAP25 using SNAP25 Sigma S9684 and Synaptic system SYSY111-111 antibodies in wildtype (WT) and SNAP25 knockout (KO) primary hippocampal cultures lysate (N=2). WT and KO primary hippocampal cultures are made from SNAP25 KO embryos (E18-19) and lentivirus containing either an empty backbone (no rescue, KO) or WT SNAP25 (WT)(*1*).  SNAP25 antibody (SYSY 111-11) is used as a control antibody of SNAP25 while B-actin (Santa Cruz sc-47778) is used as a loading control. Similar to SYSY 111-111, SNAP25 antibody (S9684) selectively recognized SNAP25 only in the wildtype (WT) primary hippocampal culture lysate.

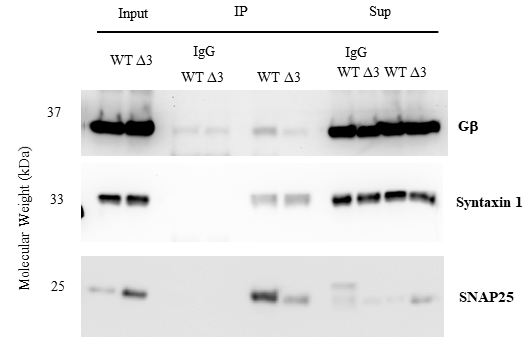

**Supplementary Figure 2. Western blot analysis of SNAP25 coIP.** To understand the SNARE complex co-immunoprecipitation (coIP) with SNAP25 antibody, we compared the components of the SNARE complex coIPed from epinephrine stimulated wildtype (WT) and SNAP25∆3 mice (∆3), truncated c-terminus SNAP25 (Δ3), causing two-fold reduction in Gβγ-SNARE interaction(*2*). We detected SNAP25 antibody specific immunoprecipitation of Gβ, Syntaxin1, and SNAP25 suggesting that we have successfully coIPed either t-SNARE or ternary SNARE.

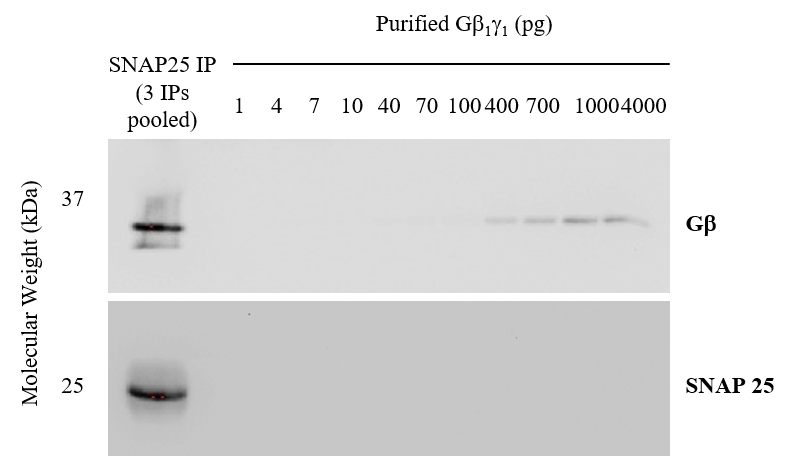

**Supplementary Figure 3. Quantification of Gβ precipitated with SNAP25.** Quantitative western blot analysis estimated approximately 1µg Gβγ was pulled down with SNAP25 per half brain.
